# MRCK controls myosin II activation in the polarized cortex of mouse oocytes and promotes spindle rotation and male pronucleus centration

**DOI:** 10.1101/2022.09.25.509421

**Authors:** Anne Bourdais, Benoit Dehapiot, Guillaume Halet

**Affiliations:** Univ Rennes, CNRS, IGDR - UMR 6290, F-35000 Rennes, France

## Abstract

Asymmetric meiotic divisions in oocytes rely on spindle positioning in close vicinity to the cortex. In mouse oocytes arrested at metaphase II, eccentric spindle positioning is associated with a chromatin-induced remodeling of the overlying cortex, including the build-up of an actin cap surrounded by a ring of activated myosin II. While the role of the actin cap in promoting polar body formation was demonstrated, the role of ring myosin II, and its mechanism of activation, have remained elusive. Here, we show that ring myosin II activation requires Myotonic dystrophy kinase-Related Cdc42-binding Kinase (MRCK), downstream of polarized Cdc42. During anaphase-II, inhibition of MRCK resulted in spindle rotation defects and a decreased rate of polar body emission. Remarkably, some oocytes eventually achieved spindle rotation by disengaging one cluster of chromatids from the anaphase spindle. We show that the MRCK/myosin II pathway also regulates the flattening of the fertilization cone to initiate male pronucleus centration. These findings provide novel insights into mammalian oocyte polarization and the role of cortical myosin II in orchestrating asymmetric division.

## Introduction

In preparation for embryonic development, oocytes must haploidize their maternal genome. In mammalian oocytes, this is orchestrated through two highly asymmetric meiotic divisions, leading to the formation of two small polar bodies in which extra chromosomes are discarded. To achieve these highly asymmetric divisions, oocytes position their meiotic spindle in close vicinity to the cortex, leading to cortical reorganization known as oocyte polarization (Chaigne et al., 2012; Yi et al., 2013; Mogessie et al., 2018). In ovulated mouse oocytes arrested at metaphase II (MII), a classical feature of cortical polarization is the establishment of a thick actin cap delineated by a stationary ring of activated nonmuscle myosin II, thereby defining the polar body-forming region (Maro et al., 1984; Longo and Chen, 1985; Van Blerkom and Bell, 1986; Deng et al., 2007; Wang et al., 2011). Remarkably, mouse oocyte polarization exemplifies a non genomic role of DNA, whereby cortical reorganization is driven from a distance, via a cytoplasmic gradient of activated Ran GTPase generated by maternal chromosomes (Deng et al., 2007; Yi et al., 2011; Dehapiot and Halet, 2013; Wang et al., 2020; Mori et al., 2021). Accordingly, the polarity pathway is conserved during anaphase II (AII), with the establishment of two smaller actin caps each surrounded by a ring of myosin II, in the cortical protrusions overlying segregated chromatids (Wang et al., 2020; Dehapiot et al., 2021).

In an effort to describe further the molecular cascade for cortical polarization, we and others previously identified the small GTPase Cdc42 as a key intermediate in the formation of the actin cap (Dehapiot et al., 2013; Dehapiot and Halet, 2013; Wang et al., 2013). As a likely downstream effector of Cdc42, the Arp2/3 complex is enriched in the polarized cortex and promotes the assembly of the actin cap, suggesting that the latter is organized as a branched network (Pollard, 2007; Yi et al., 2011; Zhang et al., 2017; Bourdais et al., 2021). Through the use of diverse inhibitors and molecular tools, the role of the actin cap in promoting polar body formation was demonstrated (Maro et al., 1984; Schatten et al., 1986; Terada et al., 2000; Dehapiot et al., 2013; Wang et al., 2013). In addition, Arp2/3 inhibition was shown to induce MII spindle drifting away from its eccentric position, due to reversed cytoplasmic streaming powered by myosin II (Yi et al., 2011). These findings point to a functional crosstalk between actin and myosin II cytoskeletal assemblies. However, the mechanism underlying ring myosin II activation, and its role during oocyte meiotic divisions, remain unclear. Intriguingly, we recently uncovered that the active myosin II ring is lost upon Cdc42 inactivation (Dehapiot et al., 2021). Therefore, while the current view is that the myosin II ring is driven by myosin light chain kinase/MLCK (Deng et al., 2005; Matson et al. 2006; Deng et al., 2007; Ajduk et al., 2011; McGinnis et al., 2015; Mackenzie et al., 2016), our observations rather support a Cdc42-dependent mechanism. In line with this view, we previously reported that Cdc42-inhibited MII oocytes do not experience spindle drift, despite a loss of the actin cap (Dehapiot et al., 2013; Dehapiot and Halet, 2013).

During the early stages of AII in fertilized oocytes, the spindle remains parallel to the cortex, but then reorients perpendicularly in a process referred to as spindle rotation, allowing for internalization of one cluster of chromatids, while the other is discarded into the second polar body (PB2) (Maro et al., 1984). Significant advances were recently made in our understanding of spindle rotation, which was shown to result from spontaneous symmetry breaking in cortical polarization signals, combined with unilateral furrow ingression serving as a driving force (Wang et al., 2020; Dehapiot et al., 2021). While actomyosin contractility as a whole is critical to PB2 formation and cytokinesis, the specific contribution of ring myosin II could not be firmly established, due to the lack of discriminating tools.

To gain insight into ring myosin II activation and function, we explored further its regulation by Cdc42. We show that ring myosin II requires the polarized activation of MRCKβ (also known as Cdc42bpb), downstream of Cdc42 activation. The MRCK/myosin II pathway is conserved during anaphase II (AII) and drives ring myosin II activation in the cortical protrusions overlying segregated chromatids. MRCK inhibition resulted in a complete loss of ring myosin II, while the cleavage furrow myosin II pool was preserved, owing to spatially segregated Cdc42 and RhoA zones. The loss of ring myosin II was associated with spindle rotation defects during AII, leading to spindle distorsion and a failure of PB2 cytokinesis. Strikingly, a fraction of oocytes eventually achieved symmetry breaking through disengaging one cluster of chromatids from the AII spindle, thus enabling rotation to proceed. In addition, we show that in fertilized oocytes, a similar Cdc42/MRCK/myosin II module promotes the flattening of the fertilization cone for inward migration of the male pronucleus.

## Results

### Polarized myosin II activation relies on Cdc42 signaling

To explore the role of Cdc42 in the establishment of the myosin II ring, MII oocytes were treated with the allosteric Cdc42 inhibitor ML-141 (Hong et al. 2013) and examined for MRLC phosphorylation at Ser19 (P-MRLC) as a proxy for myosin II activation (Vicente-Manzanares et al., 2009; Heissler and Sellers, 2016). In control oocytes treated with vehicle (DMSO), P-MRLC showed the landmark ring accumulation at the rim of the F-actin cap (Fig. 1A). In contrast, treatment with ML-141 induced a complete loss of the P-MRLC ring (Fig. 1A,G), thus recapitulating our previous observations with Cdc42T17N expression (Dehapiot et al., 2021). Cdc42 inhibition was also invariably associated with a partial or complete loss of the polarized F-actin cap (Fig. 1A, white arrow), consistent with Cdc42 acting as a key intermediate in chromatin-induced F-actin polarization. Transcriptome and translatome profiling of mouse MII oocytes identified the expression of the two predominant myosin II heavy chain paralogs, myosin-IIA and –IIB, while myosin-IIC was not detected (Tang et al., 2009; Hu et al., 2022). Accordingly, myosin-IIA distributed as a polarized ring in the cortex of MII oocytes, while myosin-IIB was not detected at that stage (Fig. S1A,B).

**Figure 1.**
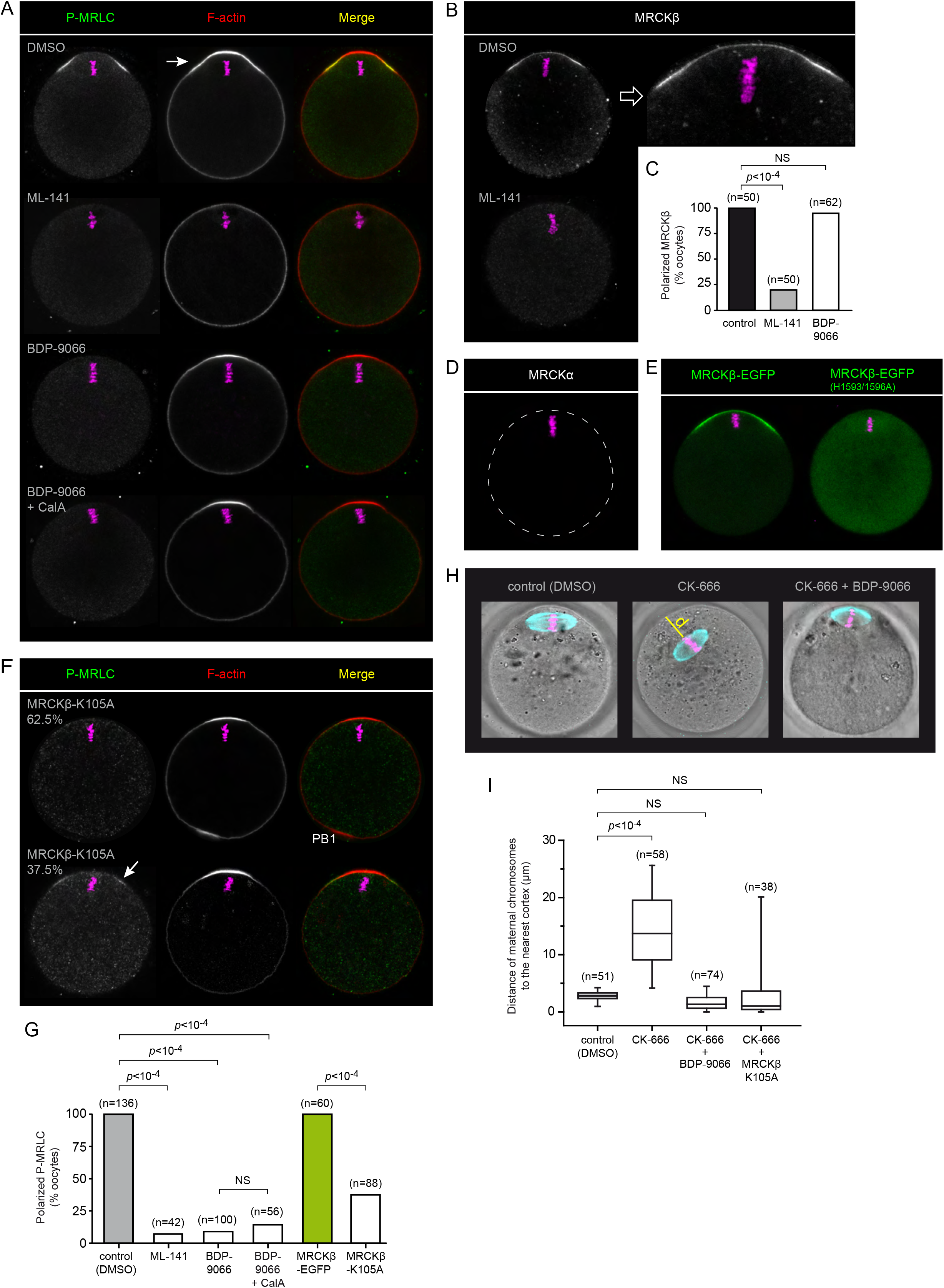
MRCK promotes myosin II activation in the polarized cortex. **(A)** Immunofluorescence detection of activated myosin II (P-MRLC) in MII oocytes treated for 1 h with DMSO, ML-141 (5 μM), BDP-9066 (1 μM) or a combination of BDP-9066 (1 μM) and calyculin A (CalA; 0.5 nM). Images are single confocal frames taken across metaphase chromosomes. **(B)** Immunofluorescence detection of endogenous MRCKbeta in MII oocytes. The top row shows a control oocyte treated with DMSO. An enlarged view of the polarized cortex is shown on the right. The bottom row shows an oocyte treated with ML-141 (5 μM). **(C)** Bar graph depicting the fraction of oocytes showing polarized accumulation of MRCKbeta, in MII oocytes treated with DMSO, ML-141 or BDP-9066. *P* values were calculated using Fisher’s exact test. **(D)** Immunofluorescence detection of MRCKα in a mouse MII oocyte. The dotted line denotes the oocyte perimeter. No cortical staining could be detected. The image is representative of 33 similar observations. **(E)** Localization of MRCKβ-EGFP (left) and MRCKβ H1593/1596A-EGFP (right) in MII oocytes. Note the absence of cortical localization for the Cdc42 binding-deficient MRCKβ. Images are representative of over 40 similar observations. **(F)** Immunofluorescence detection of activated myosin II (P-MRLC) in MII oocytes expressing MRCKβ-K105A. A majority of oocytes showed a complete loss of the P-MRLC ring (62.5%; top row), while the remaining oocytes showed incomplete inhibition (37.5%; bottom row, white arrow). **(G)** Bar graph depicting the percentage of MII oocytes showing a P-MRLC ring, in various experimental conditions as shown in (A), (E) and (F). *P* values were calculated using Fisher’s exact test. **(H)** Immunofluorescence detection of tubulin showing spindle localization in MII oocytes treated with DMSO (left), CK-666 (100 μM) for 3h (middle), and CK-666 (100 μM) and BDP-9066 (1 μM) for 3h (right). The yellow line shows the distance (d) separating the maternal chromosomes from the nearest cortex. **(I)** Box plot showing the distance between maternal chromosomes and the nearest cortical region in oocytes treated with DMSO (controls), CK-666 alone, CK-666 and BDP-9066, and in oocytes expressing MRCKβ-K105A treated with CK-666. *P* values were calculated using Student’s *t*-test. NS: non significant. F-actin was labeled with Alexa Fluor 568-phalloidin. DNA was stained with TO-PRO-3. The number of oocytes scored is indicated above each bar/boc (C,G,I). Scale bars represent 10 μm.

Based on the use of compound ML-7, previous studies have argued that the myosin II ring was driven by MLCK, a kinase primarily activated by Ca^2+^/calmodulin (Hong et al., 2011). However, we could not achieve significant myosin II inhibition using 15 μM ML-7 (Fig. S2). To clarify further the contribution of MLCK, we employed three alternative inhibitory strategies: (1) Ca^2+^ depletion (Miao et al., 2012), (2) injection of peptide-18 (Lukas et al., 1999), and (3) treatment with 1 μM wortmannin, a potent and irreversible MLCK inhibitor (Nakanishi et al., 1992; Davies et al., 2000). However, none of these treatments demonstrated any inhibitory effect on the P-MRLC ring (Fig. S2), arguing strongly against MLCK as the MRLC kinase.

Another two classic MRLC kinases are ROCK and group I PAKs, which act downstream of RhoA and Cdc42/Rac, respectively (Amano et al., 1996; Chew et al., 1998; Wilkinson et al., 2005). A role for ROCK appears highly unlikely since RhoA and ROCK are absent from the polarized cortex, and the P-MRLC ring is resistant to ROCK inhibition (Dehapiot et al., 2021). Likewise, IPA-3, an allosteric inhibitor of group I PAKs (Deacon et al., 2008), did not prevent polarized myosin II activation (Fig. S2). Collectively, these data suggest that polarized myosin II localization and activation in MII oocytes involve a kinase acting downstream of Cdc42, other than PAK.

### Cdc42 recruits MRCKβ to the polarized cortex for myosin II activation

MRCK is a conserved serine/threonine kinase which promotes myosin II activation directly via MRLC Ser19 phosphorylation (Leung et al., 1998; Tan et al., 2011). Owing to a canonical Cdc42/Rac Interactive Binding (CRIB) domain located at its C terminus (Fig. S3), MRCK is targeted to sites of Cdc42 activation, making it a valuable candidate for polarized myosin II activation (Ando et al., 2013; Unbekandt and Olson, 2014; Zhao and Manser, 2015). Consistent with this idea, MRCKβ was detected in the polarized cortex of MII oocytes (Fig. 1B,C). In contrast, the closely related MRCKα was not detected (Fig. 1D). Consistent with Cdc42 acting as a polarizing cue, treatment with ML-141 inhibited the cortical localization of MRCKβ (Fig. 1B,C). Similarly, a MRCKβ-EGFP construct (Ando et al., 2013) decorated the polarized cortex, while a mutant MRCKβ impaired for Cdc42·GTP binding (MRCKbeta H1593/1596A; Leung et al., 1998; Ando et al., 2013), remained cytosolic (Fig. 1E and S3).

We next examined the contribution of MRCK to the polarized activation of myosin II. Because genetic ablation of MRCK was reported to substantially rewire kinase signaling, and may preclude kinase-independent functions (Kurimchak et al., 2020), we first aimed for a dominant-negative approach (Leung et al., 1998; Nakamura et al., 2000; Tan et al., 2008). Thus, a kinase-dead MRCKβ mutant was generated by substituting lysine 105 in the kinase domain, for alanine (K105A) (Unbekandt et al., 2020). Accordingly, expression of MRCKβ-K105A via cRNA injection abolished the P-MRLC ring in a majority (62.5%) of injected oocytes, while the expression of MRCKβ-EGFP had no effect (Fig. 1F,G). The remaining oocytes (37.5%) showed residual P-MRLC signal (Fig. 1F,G), likely reflecting lower expression of the kinase-dead mutant. Alternatively, we challenged oocytes with the recently characterized small molecule inhibitor BDP9066, which is highly selective for MRCK, while having no effect on MLCK (Unbekandt et al., 2018). Remarkably, acute MRCK inhibition with BDP9066 (1 μM, 1h) abolished MRLC phosphorylation in virtually all oocytes examined (Fig. 1A,G). The myosin-IIA ring was lost accordingly, consistent with disassembly of the actomyosin filaments (Fig. S1A).

MRCK may also activate myosin II indirectly, via the phosphorylation of the protein phosphatase 1 (PP1) regulatory subunit MYPT1, leading to inactivation of myosin phosphatase activity (Tan et al., 2001; Wilkinson et al., 2005). However, supplementation of the culture medium with calyculin A, a potent PP1 inhibitor (Ishihara et al., 1989), did not rescue ring P-MRLC in MRCK-inhibited oocytes (Fig. 1A,G). While this observation does not rule out MYPT1 as an additional substrate, it supports the view that oocyte MRCK acts directly as the MRLC S19 kinase. Collectively, the above data indicate that Cdc42 activation acts as a spatial cue for MRCKβ recruitment at the polarized cortex, leading to MRLC phosphorylation and establishment of the active myosin II ring.

To establish the functional inhibition of ring myosin II in MRCK-inhibited oocytes, we employed CK-666 to inhibit the Arp2/3 complex and induce MII spindle drift powered by myosin II contractility (Nolen et al., 2009; Yi et al., 2011). Thus, oocytes treated with CK-666 (100 μM; 3h) alone demonstrated a substantial drift of the spindle toward the cell interior (Fig. 1H,I). In contrast, the spindle remained closely apposed to the cortex in oocytes treated with both CK-666 and BDP9066 (Fig. 1H,I). In addition, we noticed that inhibition of ring myosin II was associated with oocyte rounding, through the loss of the outward bulge which otherwise defines the amicrovillar polarized cortex (Fig. 1A). These results are consistent with a loss of actomyosin contractility in the polarized cortex of MRCK-inhibited oocytes. Importantly, these data also demonstrate that inactivation of ring myosin II is not associated with a drift of the MII spindle away from the cortex, contrary to previous studies using ML-7 (Deng et al., 2005; Deng et al., 2007; McGinnis et al., 2015).

### Crosstalk between ring myosin II and the Arp2/3 complex shapes the polarized cortex

Ring myosin II and the Arp2/3 complex were recently suggested to engage into a mutual inhibition at the polarized cortex, whereby myosin would restrict the size of the actin cap (Wang et al., 2020). However, this conclusion was reached via artificial activation of Rho at the cortex, which may prove antagonistic toward Cdc42 signaling (Vaughan et al., 2011). We performed MRCK inhibition to test whether ring myosin II opposes Arp2/3. Contrary to this prediction, MRCK-inhibited oocytes showed a significant reduction of the size of the actin cap, though we never observed a complete loss of the cap (Fig. 2A). Interestingly, we uncovered that the outer region of the polarized Arp2/3 domain overlaps with ring myosin II (Fig. 2B, white arrows). Moreover, MRCK inhibition resulted in a loss of the outer region of the Arp2/3 cap, matching the loss of the P-MRLC ring (Fig. 2B). These data reveal an unanticipated role of ring myosin II in the broadening of the polarized Arp2/3 domain, and hence the actin cap.

**Figure 2.**
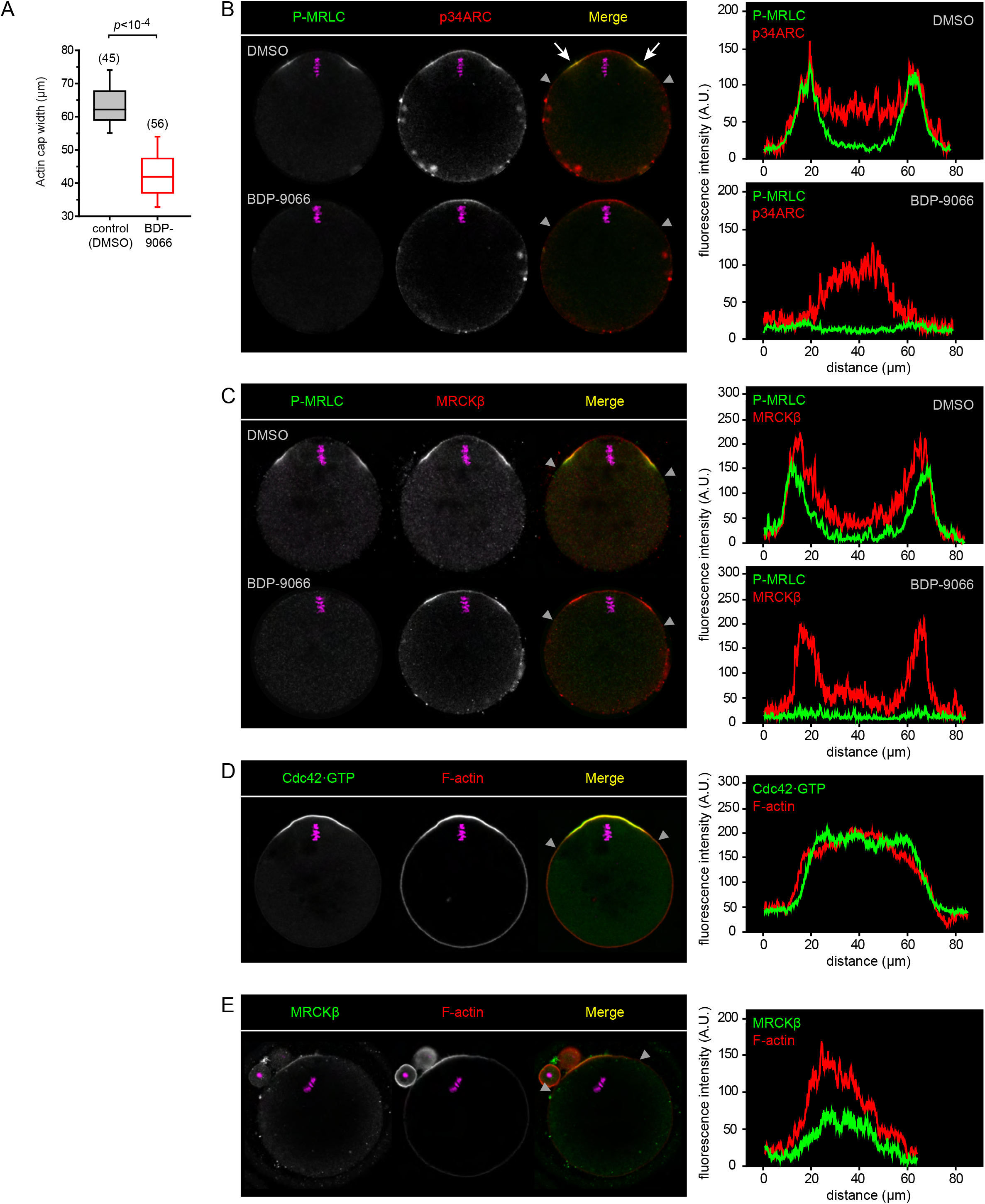
MRCKβ co-localizes with ring myosin II and Arp2/3 at the polarized cortex. **(A)** Box plot showing the width of the actin cap in control MII oocytes treated with DMSO and MII oocytes treated with BDP-9066 (1 μM). *P* value was calculated using Student’s *t*-test. The number of oocytes scored is indicated above each box. **(B)** Immunofluorescence detection of activated myosin II (P-MRLC) and the Arp2/3 complex (p34ARC) in MII oocytes treated for 1 h with DMSO (top row) or BDP-9066 (1 μM; bottom row). White arrows point to the co-localization of P-MRLC and p34ARC at the shoulders of the actin cap. **(C)** Immunofluorescence detection of activated myosin II (P-MRLC) and MRCKβ in MII oocytes treated for 1 h with DMSO (top row) or BDP-9066 (1 μM; bottom row). **(D)** Detection of Cdc42·GTP in a MII oocyte expressing the Cdc42 biosensor MCB-EGFP. **(E)** Immunofluorescence detection of MRCKβ and F-actin in MII oocytes treated for 3 h with 100 μM CK-666. In (B-E), fluorescence intensity profiles are shown, corresponding to the polarized cortex region delinated by the two grey arrowheads in the merge images. F-actin was labeled with Alexa Fluor 568-phalloidin. DNA was stained with TO-PRO-3. Scale bars represent 10 μm.

MRCKβ demonstrated comparable accumulation at the rim of the F-actin cap, matching the P-MRLC ring (Fig. 2C). This ring enrichment was unaffected by myosin II inhibition (Fig. 2C), ruling out a mechanotransduction mechanism (Kuo et al., 2011; Munjal et al., 2015). We therefore asked whether the anchoring GTPase Cdc42 was also enriched as a ring. To test this hypothesis, we designed a Cdc42·GTP biosensor derived from the minimal Cdc42·GTP-binding fragment of human MRCKβ, that we referred to as <u>M</u>RCK-derived <u>C</u>dc42·GTP <u>b</u>iosensor (MCB; Fig. S3A,B). Accordingly, MCB-EGFP decorated the polarized cortex of MII oocytes, and was released to the cytosol upon treatment with ML-141 (Fig. S3C,D). Likewise, substitution of the two critical histidine residues in the CRIB domain (H1593/1596A; Leung et al., 1998; Ando et al., 2013) abolished cortical localization (Fig. S3E). Fluorescence profile analysis revealed that the Cdc42·GTP signal was maximal in the central region of the polarized cortex and faded gradually with increasing distance, much similar to the F-actin cap signal (Fig. 2D). While this pattern likely reflects upstream regulation by the Ran·GTP gradient, it is inconsistent with MRCKβ clustering as a ring. In contrast, inactivation of the Arp2/3 complex induced a redistribution of MRCKβ into a polarized cap, as previously described for ring myosin II (Yi et al., 2011). Intriguingly, the bulk of the actin cap was still present, in line with the fact that CK-666 does not disassemble preformed branches (Hetrick et al., 2013), and arguing for a fairly slow turnover of actin filaments in the cap. This observation suggest that the ring pattern is not due to the passive exclusion of MRCK/myosin II from the branched network. Rather, the circumferential actin flow generated by polarized Arp2/3 activation, may transport MRCK/myosin II for clustering at the rim of the actin cap, analogous to actin flow-mediated myosin-IIA partitioning in polarized fibroblasts (Yi et al., 2011; Beach et al., 2017).

### Cdc42 and RhoA zones define two cortical myosin II pools in anaphase-II

We recently uncovered that upon oocyte activation, the P-MRLC ring splits into two smaller rings overlying the two clusters of segregated chromatids, while a second pool of myosin II is activated de novo by the RhoA/ROCK pathway to drive unilateral membrane furrowing above the central spindle (Dehapiot et al., 2021). We therefore inferred that the Cdc42/MRCK pathway remains active in anaphase-II, yet spatially segregated from the RhoA/ROCK pathway. To substantiate this view, we examined the spatial distribution of activated Cdc42 and RhoA GTPases and their cognate effector kinases. In live MII oocytes, Cdc42·GTP was readily detected in the polarized cortex, while RhoA activation, monitored using the mcherry-AHPH biosensor (Piekny et al., 2008), remained undetectable (Fig. 3A). Following activation with SrCl2 to trigger cell cycle progression to anaphase II, the Cdc42·GTP cap split into two smaller caps associated with each of the two chromatid clusters, while the intervening cortical region showed RhoA·GTP accumulation (Fig. 3A). Accordingly, MRCKβ localized selectively in the two cortical protrusions overlying the chromatid clusters, while ROCK1 concentrated in a narrow region over the central spindle (Fig. 3B,C). Myosin-IIA was detected both in the rings and in the ingressing furrow (Fig. S1C), in line with previous observations (Sharif et al., 2015). Of note, myosin-IIB became detectable at the cortex of anaphase-II oocytes, and concentrated exclusively in the cytokinetic furrow (Fig. S1D).

**Figure 3.**
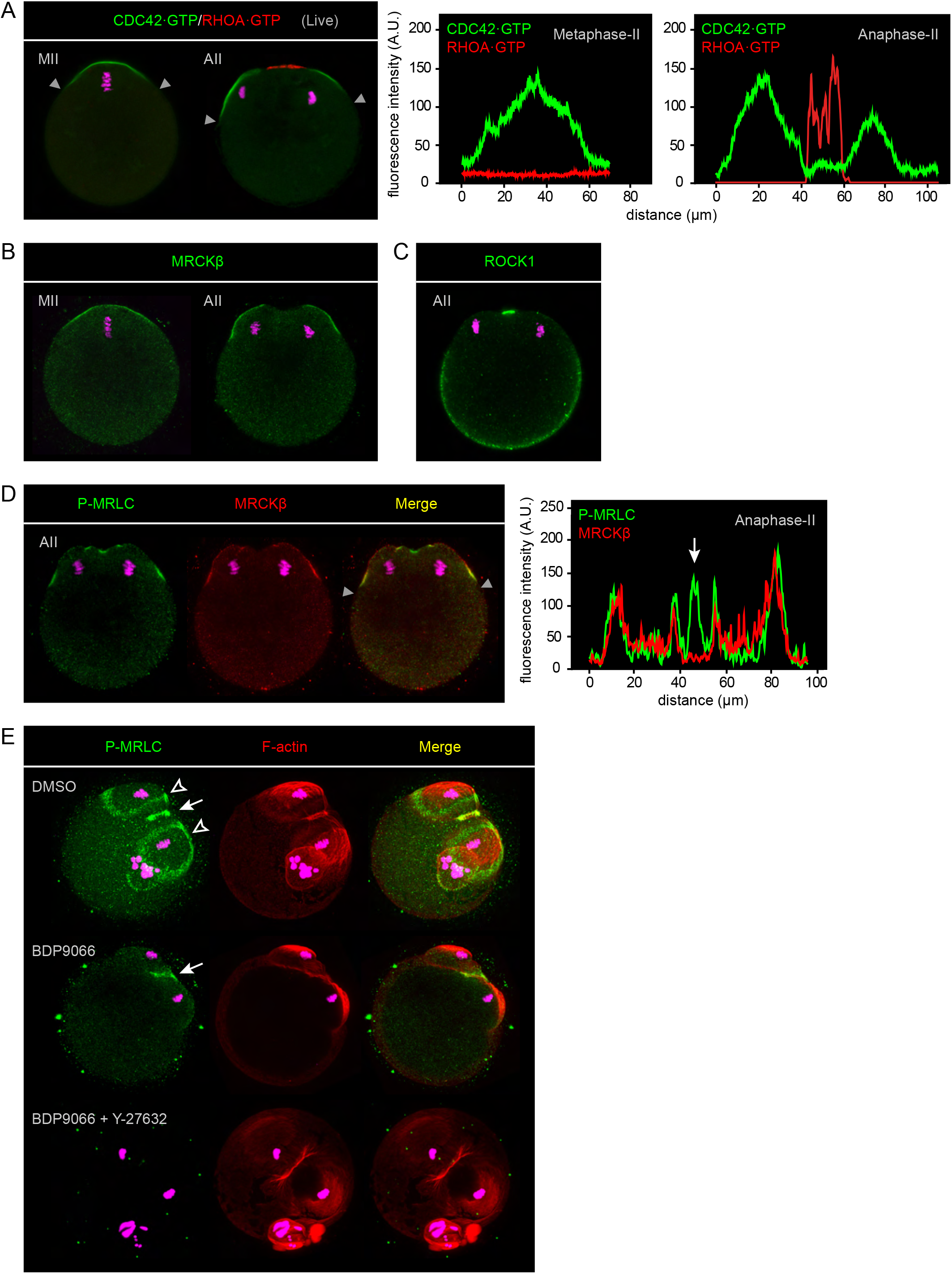
Two distinct myosin II pools coexist in the cortex of activated oocytes. **(A)** Detection of Cdc42·GTP (green) and RhoA·GTP (red) in live oocytes at the metaphase-II (MII, left) and anaphase-II (AII, right) stages. Oocytes were injected at the MII stage with cRNAs encoding the MCB-EGFP and mcherry-AHPH biosensors. Images are Z-compressions of 3 consecutive confocal frames. **(B)** Immunofluorescence detection of MRCKβ in fixed MII (left) and AII (right) oocytes. **(C)** Immunofluorescence detection of ROCK1 in an AII oocyte. **(D)** Immunofluorescence detection of activated myosin II (P-MRLC) and MRCKβ in an AII oocyte. **(E)** Immunofluorescence detection of activated myosin II (P-MRLC) in the cortex of activated oocytes undergoing anaphase II. Oocytes were treated with DMSO (top row), BDP-9066 (middle row) or a combination of BDP-9066 and Y-27632 (bottom row). Open arrowheads point to the P-MRLC rings overlying the chromatid clusters in the control (DMSO) oocyte. White arrows point to cytokinetic P-MRLC in the control (DMSO) and BDP-9066-treated oocytes. Images are z-compression of 33 (DMSO), 15 (BDP-9066) and 30 (BDP-9066 + Y-27632) consecutive confocal frames. F-actin was labeled with Alexa Fluor 568-phalloidin. DNA was stained with SiR-DNA (panel A) or TO-PRO-3 (panels BE). Fluorescence intensity profiles correspond to the cortical regions delineated by the grey arrowheads in the merge images. In (D), the white arrow points to the ingressing furrow region that is enriched in P-MRLC but devoided of MRCKβ.

Consistent with our observations in MII oocytes, MRCKβ co-localized precisely with the two smaller P-MRLC rings, but not with cytokinetic P-MRLC in the cleavage furrow (Fig. 3D). Accordingly, MRCK inhibition resulted in the selective loss of ring P-MRLC, while cytokinetic P-MRLC was preserved (Fig. 3E). Furthermore, simultaneous treatment with MRCK (BDP9066) and ROCK (Y-27632) inhibitors resulted in a complete loss of cortical myosin II activation (Fig. 3E).

Together with our previous findings on the activation of the RhoA/ROCK pathway during AII (Dehapiot et al., 2021), the above data provide strong evidence for the activation of two distinct myosin II populations in the cortex of activated oocytes, downstream of spatially segregated Cdc42/MRCK and RhoA/ROCK pathways. While the role of the ROCK/myosin II module in driving cytokinetic furrow ingression has been demonstrated, the role of ring myosin II in activated oocytes has remained elusive (Wang et al., 2020; Dehapiot et al., 2021). We therefore performed MRCK inhibition with BDP9066 to clarify the role of ring myosin II during anaphase II.

### MRCK promotes spindle rotation in activated oocytes

To find out what role ring myosin II might play during oocyte activation, we performed live imaging of spindle and chromosome dynamics in SrCl2-activated oocytes. As previously reported (Maro et al., 1984; Wang et al., 2020; Dehapiot et al., 2021), vehicle (DMSO)-treated control oocytes initiated anaphase II (AII) in a symmetrical fashion, whereby the AII spindle lied parallel to the cortex, leading to the bulging of two membrane protrusions over each set of segregated chromatids (Fig. 4A; video 1). This configuration was short-lasted as unilateral furrowing above the spindle equator was quickly followed by spindle rotation, culminating in the emission of the PB2, while a single haploid set of maternal chromosomes was retained in the oocyte (Fig. 4A,D; video 1). During the course of rotation, the internalized cluster of chromatids remained firmly connected to the anaphase spindle, and was thus relocated deeper into the inner cytoplasm, while the overlying cortical protrusion resolved (video 1). Accordingly, upon meiotic exit, a single female pronucleus formed in the oocyte cytoplasm, at a distance from the cortex (Fig. 4E, left).

**Figure 4.**
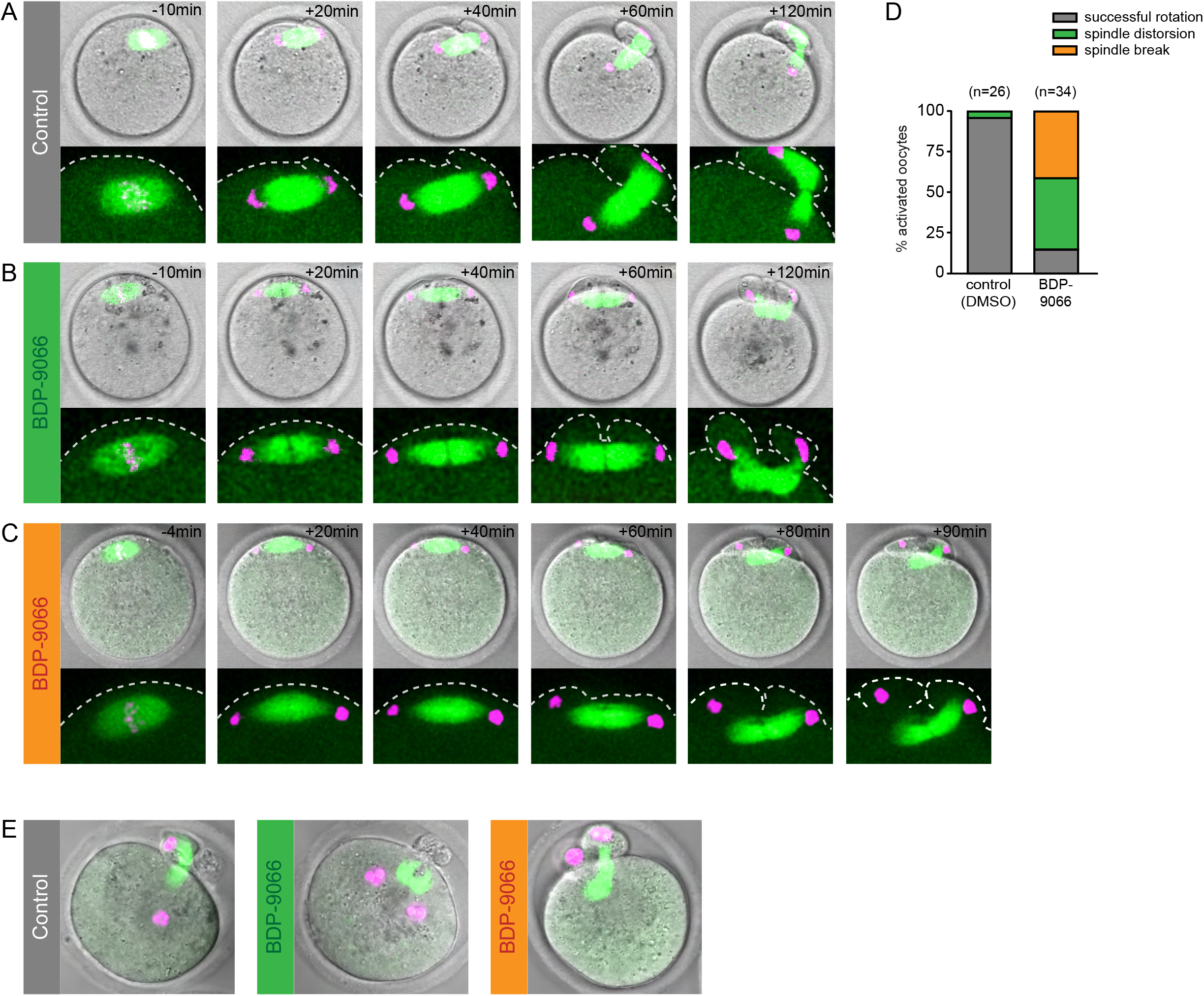
MRCK promotes spindle rotation in anaphase-II. **(A)** Live imaging of a SrCl2-activated MII oocyte, treated with DMSO as a control. Still images are from Video 1. **(B)** Live imaging of a SrCl2-activated MII oocyte, treated with BDP9066. Still images are from Video 2. **(C)** Live imaging of a SrCl2-activated MII oocyte, treated with BDP9066. Still images are from Video 3. Labels indicate the time to, or from, the onset of chromatid segregation. **(D)** Bar graph depicting the percentage of activated oocytes showing normal spindle rotation, the spindle distorsion phenotype, or the spindle break phenotype. The total number of oocytes scored is indicated in parentheses. **(E)** Representative images of pronucleus configuration in activated oocytes imaged live 6 h post-insemination. Before activation, oocytes were injected with cRNA encoding EGFP-MAP4 to label microtubules. Chromosomes were labeled with H2B-mcherry.

Oocytes treated with the MRCK inhibitor BDP9066 initiated anaphase II with a similar time course as controls (Fig. 4B,C). However, while a small fraction of these oocytes achieved symmetry breaking to form a PB2 (Fig. 4D), the remaining oocytes exhibited striking spindle rotation defects. Thus, at the onset of furrow ingression, the spindle remained parallel to the cortex for an extended duration, with both chromatid clusters closely apposed to the cortex. In about half of these oocytes, we observed a severe distorsion of the AII spindle while the cleavage furrow ingressed further, as neither of the two chromatid clusters would let go of cortical anchoring. This lead to the bulging of two polar body-like protrusions, each containing one set of chromatids (Fig. 4B,D; video 2). This distorted configuration persisted until meiotic exit, leading to the formation of binucleate parthenotes (Fig. 4E, middle). In the remaining MRCK-inhibited oocytes, we observed a striking phenomenon whereby one of the membrane-apposed chromatin clusters disengaged from the AII spindle pole at the onset of cortical ingression (Fig. 4C,D; video 3). As furrowing progressed further, spindle rotation occurred in a consistent fashion, with the chromatin-associated spindle pole forming the PB2, while the chromatin-less spindle pole reoriented toward the cell interior (Fig. 4C; video 3). Remarkably, the detached cluster remained in close vicinity to the overlying cortex, and formed a pronucleus that was often still enclosed in the polar body-like protrusion at the time of fixation (Fig. 4E, right).

Collectively, these data suggest that MRCK is required for normal spindle rotation in oocytes undergoing anaphase-II. While membrane furrowing above the central spindle appears unaffected, the major defect of MRCK-inhibited oocytes appears to lie in the inability of chromatid clusters to let go of their cortical anchoring, thus precluding spindle rotation.

### MRCK promotes myosin II activation in the fertilization cone and centration of the male pronucleus

Shortly after sperm fusion with the oocyte, the so-called fertilization cone (FC) is formed at the site of sperm entry. The general features of the FC are reminiscent of the polarized cortex overlying the MII spindle, i.e. an amicrovillar membrane protrusion enriched in F-actin and myosin II, which formation requires nearby chromatin signaling (Maro et al., 1984; Simerly et al., 1998; Deng et al., 2005; Deng and Li, 2009; Ajduk et al., 2011). In the same line, we observed that active myosin II (P-MRLC) localized as a ring at the boundary of the FC (Fig. 5A,B and S4A), akin to the P-MRLC ring overlying maternal chromosomes. In addition, activated myosin II similarly overlapped with the rim of the FC actin cap (Fig. 5A). We therefore surmised that MRCK may similarly regulate myosin II activation in the FC. Consistent with this idea, the FC was enriched in Cdc42·GTP (Fig. S4B), and MRCKβ accumulated as a ring at the boundary of the FC, matching the P-MRLC ring (Fig S4C,D). Significantly, supplementation of the fertilization medium with BDP9066 (1 μM) abolished the FC P-MRLC ring, and shortened the FC actin cap accordingly (Fig. 5A,B). These data suggest that myosin II activation in the FC also relies on the Cdc42/MRCK pathway.

**Figure 5.**
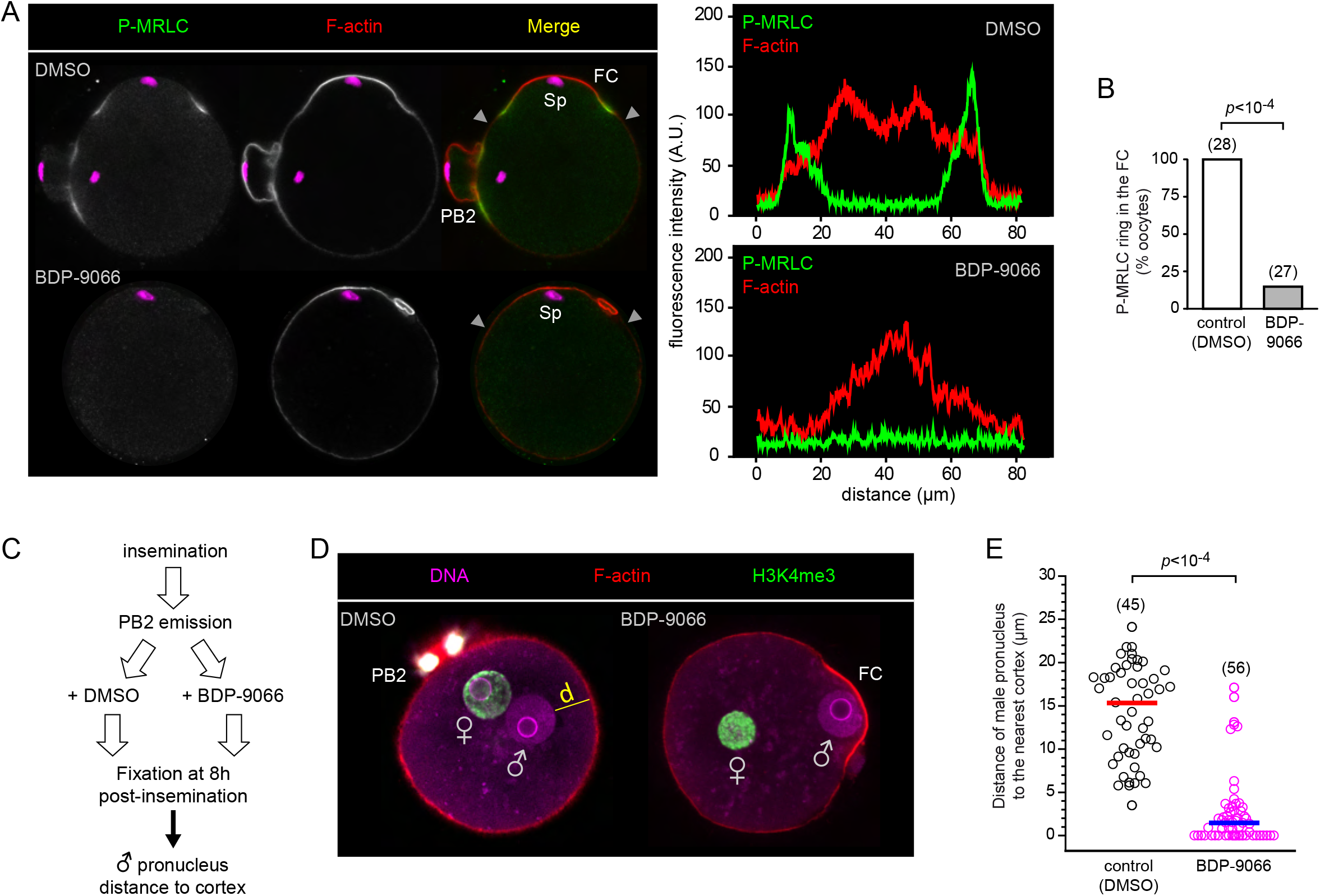
MRCK promotes myosin II activation in the fertilization cone and male pronucleus centration. **(A)** Immunofluorescence detection of activated myosin II (P-MRLC) in fertilized oocytes. Top row: control oocyte treated with DMSO. Bottom row: oocyte fertilized in the presence of BDP-9066. Note the P-MRLC ring at the base of the fertilization cone (FC) in the control oocyte, and absence of in the MRCK-inhibited oocyte. Oocytes were fixed 3 hours postinsemination. PB2: second polar body. Sp: sperm chromatin. Fluorescence intensity profiles correspond to the cortical regions delineated by the grey arrowheads in the merge images. **(B)** Bar graph depicting the percentage of fertilized oocytes showing a P-MRLC ring at the base of the FC. The fertilization medium was supplemented with DMSO (control) or BDP-9066. *P* value was calculated using Fisher’s exact test. **(C)** Schematic illustrating the experimental protocol for monitoring male pronucleus migration. **(D)** Detection of male and female pronuclei in zygotes fixed 8h after insemination. The left image shows a control zygote treated with DMSO after PB2 emission. The yellow line denotes the distance (d) of the male pronucleus to the nearest cortex. The right image shows a zygote treated with BDP-9066 after PB2 emission. Note the expanded male pronucleus still apposed to the FC cortex. Images are z-compressions of two consecutive confocal frames. **(E)** Scatter plot of the male pronucleus distance to the nearest cortex, as depicted in (D). Red and blue horizontal bars show the median values. *P* value was calculated using a two-tailed Mann-Whitney U test. DNA was labeled with TO-PRO-3. F-actin was labeled with Alexa Fluor 568-phalloidin. Female pronuclei were immuno-stained for H3K4me3 (green).

Previous studies have demonstrated that the protrusion of a FC is not a prerequisite for successful sperm fusion and oocyte activation (Maro et al., 1984; Schatten et al., 1986; Terada et al., 2000). Yet, a recent study suggested that the flattening of the FC shortly after PB2 emission initiates the migration of the male pronucleus (PN) toward the center of the zygote, for apposition with the female PN ahead of the first embryonic cleavage (Scheffler et al., 2021). To investigate any defect in male PN migration, oocytes were fertilized in vitro and zygotes showing a PB2 were further cultured in the presence of DMSO (control) or BDP9066 (Fig. 5C). Zygotes were fixed 8 h post insemination, at which time the pronuclei are expected to have congressed to the center of the cell (Maro et al., 1984). As expected, control zygotes exhibited closely apposed expanded male and female PNs located near the center of the zygote (Fig. 5D,E). The male PN was unambiguously identified as being negative for H3K4me3 staining (Lepikhov and Walter, 2004). Notably, the male PN was located at a distance from the cortex, demonstrating successful migration away from the sperm entry site (Fig. 5D,E). In striking contrast, male PNs demonstrated a complete failure of migration in MRCK-inhibited oocytes, and remained apposed to an actin-rich remnant of the FC (Fig. 5D,E). These data suggest that MRCK-driven FC myosin II is required to achieve FC flattening and produce the inward directed forces that will initiate male PN migration.

## Discussion

Despite being a signature of mouse oocyte polarization, the mechanism and rationale for ring myosin II activation have been little investigated. The myosin II ring was first thought to reflect a preassembled – yet unproductive - actomyosin ring for PB2 cytokinesis (Deng et al., 2007; Yi and Li, 2012). However, we and others have shown that the driving force for unilateral membrane furrowing and PB2 formation is the RhoA-dependent activation of the ROCK/myosin II pathway above the central spindle (Zhong et al., 2005; Dehapiot et al., 2021). A major obstacle has been the lack of tools to inhibit ring myosin II specifically, without affecting ROCK/myosin II - as would do general myosin II inhibitors such as blebbistatin. This study provides a solution to this issue, as we identify MRCK as the myosin light chain-kinase acting downstream of Cdc42 to specifically activate ring myosin II, in both metaphase-II and anaphase-II.

Our data challenge the prevailing view that ring myosin II activation is driven by MLCK, based on the use of compound ML-7. Though ML-7 has not been formally tested against MRCK, it seems plausible that high concentrations may impact MRCK activity. In this regard, apical constriction of endoderm precursor cells in C. elegans gastrulation was initially reported to rely on MLCK for myosin II activation, based on the use of ML-7, but was later found to actually rely on MRCK (Lee and Goldstein, 2003; Marston et al., 2016). In mouse oocytes, conflicting results have been reported as to whether ML-7 inhibits PB2 emission (Matson et al., 2006; Larson et al., 2010; Ajduk et al., 2011; Wang et al., 2011; McGinnis et al., 2015). Moreover, oocyte-specific invalidation of the *Mylk1* gene, which encodes smooth muscle and nonmuscle MLCK, suggested that MLCK is dispensable for oocyte maturation, fertilization and polar body emission (Liang et al. 2015). These observations call for caution regarding previously reported roles for MLCK in mouse oocytes.

The loss of ring myosin II in MII oocytes did not appear to alter their overall polarized organization, aside from a flattening of the polarized cortex. While this flattening is expected to increase the polarizing signal in response to the Ran·GTP gradient (Deng et al., 2007), we observed instead a loss of the outer rim of the actin cap, suggesting that ring myosin II may stabilize and /or bundle actin filaments located in the transition zone between the actin cap and the rest of the cortex. It is unclear at this stage whether ring myosin II plays other functional roles during MII arrest, or during oocyte maturation and PB1 emission. In view of its distinctive positioning at the border of the actin cap, the myosin II ring could perhaps contribute to establishing an exclusion barrier, in order to segregate proteins that belong to the polarized cortex from those which don’t, such as sperm-binding proteins (Mori et al., 2021).

Activated oocytes in contrast, exhibited most intriguing spindle rotation defects in the absence of ring myosin II. The dramatic spindle bending and breaking phenotypes are highly suggestive of a tug-of-war between furrow ingression, which exerts a pushing force on the AII spindle (Dehapiot et al., 2021), and the chromatin clusters, which seem virtually anchored to the cortex. Most strikingky, this tug-of-war even culminated in the anaphase spindle parting from one chromatid cluster to rescue rotation. Hence, these data reveal that spindle rotation, and PB2 extrusion, can be achieved in oocytes lacking ring myosin II, providing unambiguous evidence that the myosin II ring is not the precursor of the cytokinetic ring. These findings therefore substantiate our model of spindle rotation as resulting from ROCK-induced cortical ingression, combined with chromatin-induced cortical anchoring at one pole of the anaphase spindle, independently of ring mysoin II (Dehapiot et al., 2021). Our data therefore argue against an instrumental role for asymmetric distribution of myosin II in driving spindle rotation, as was previously suggested (Wang et al., 2020). The inward-directed hydrodynamic forces detected in the region of the collapsing protrusion, as revealed by the cytoplasmic streaming pattern (Wang et al., 2020; Dehapiot et al., 2021), are more likely to arise from diminished Arp2/3 activity and the reorganization of myosin II into a cap, i.e. as a consequence of spindle rotation and chromatin distancing, not its cause. We propose that the observed spindle rotation defects arose from chromatin clusters being reluctant to let go of their cortical anchoring. Thus, ring myosin II activation may be required to prevent chromatid cluster from reaching to close to the cortex during anaphase-II. Further experiments are required to verify this assumption.

Since its discovery twenty-five years ago in drosophila, evidence for MRCK regulating actomyosin contractility has been scarce compared to the wealth of litterature on myosin II activation by ROCK (Unbekandt and Olson, 2014; Zhao and Manser, 2015). MRCK has since emerged as an important regulator of actomyosin contractility in epithelia homeostasis, cell adhesion and phagocytosis (Ando et al., 2013; Marston et al., 2016; Gagliardi et al., 2017; Zihni et al., 2017; Zihni et al., 2022). MRCK was also implicated in the regulation of cell motility and invasiveness, and has emerged as a druggable target in several cancers (Tan et al., 2008; Unbekandt et al., 2018; Kurimchak et al., 2020; East and Asquith, 2021). Here we show that MRCK is a new player in mammalian oocyte polarization and asymmetric division, downstream of polarized Cdc42. There is still much to discover on myosin II roles and regulation at the oocyte cortex. In particular, the precise architecture of actin filaments in the actin cap and FC remains to be clarified, and how this architecture is remodelled by ring myosin-II remains to be investigated.

## Materials and Methods

### Mice

All animal procedures were conducted in accordance with the European directive for the use and care of laboratory animals (2010/63/EU), and approved by the local animal ethics committee under the French Ministry of Higher Education, Research and Innovation (Project licence APAFIS#11761-2017101200282520). Mice of the MF1 strain were initially purchased from Envigo (Gannat, France) and maintained as a colony in the local animal facility.

### Oocyte recovery and culture

To minimize the number of animals, female mice (6-8 week old) were primed by intraperitoneal injection of 5-7 units of PMSG (Chronogest, MSD), followed 48h later by 5 IU hCG (Chorulon, MSD). Metaphase-II oocytes were recovered from the oviducts in M2 medium (Sigma) supplemented with 3mg/ml hyaluronidase (Sigma), followed by wash. Oocytes were subsequently cultured in home-made M16 medium (Nagy et al., 2003), in an incubator providing an atmosphere of 5% CO2 in air. To achieve Ca^2+^ depletion, MII oocytes were cultured for 1 h in Ca^2+^-free M16 medium supplemented with 1 mM EGTA, and containing 10 μM thapsigargin (Tocris) to promote intracellular Ca^2+^ store depletion (Kline and Kline, 1992a; Miao et al., 2012). To induce artificial resumption of meiosis-II, MII oocytes were cultured in Ca^2+^-free M16 medium supplemented with 10 mM SrCl2 (Kline and kline, 1992b).

### Inhibitor treatments

In our preliminary experiments, we consistently noticed that oocytes cultured in media containing bovine serum albumin (BSA) required higher concentrations of inhibitors to obtain a response similar to oocytes cultured in media in which BSA was substituted for by poly(vinyl alcohol) (PVA). Therefore, all experiments involving inhibitor treatments were performed in home-made BSA-free M16 medium supplemented with 0.05% PVA. The following small molecule inhibitors were used : ML-141 (5μM; Tocris), ML-7 (15 μM; Tocris), Wortmannin (1 μM; Echelon biosciences), IPA-3 (1 μM; Tocris), BDP-9066 (1 μM; Aobious), CK-666 (100 μM; Sigma), Y-27632 (50 μM; Merck Millipore). An equivalent amount of DMSO was used for controls. Peptide-18 (Merck Millipore) was recovered in water at a stock concentration of 1 mg/ml and microinjected in the oocyte cytoplasm.

### In vitro fertilization

Fertilization of MII oocytes was performed according to previously published protocols (Maro et al., 1984; Miao et al., 2012). Briefly, sperm recovered from 12-15 week old male MF1 mice was capacitated for 2 h in a 500-μl drop of HTF medium (Sigma), layered by mineral oil (Sigma), at 37°C in a 5% CO2 incubator. Cumulus masses were recovered from the oviducts 13 h post-hCG and were placed in a culture dish containing 2400 μl HTF, to which 50 μl of the capacitated sperm suspension was added. The dish was returned to the 5% CO2 incubator for fertilization to proceed. After 3 h, oocytes were placed in a new dish of equilibrated HTF medium to wash out unbound sperm. Fertilized oocytes were recovered at various times post-insemination for further analysis. To facilitate the fertilization of oocytes that had been micro-injected with the cRNA encoding the Cdc42·GTP biosensor, the zona pellucida was removed by a brief incubation in acidic Tyrode’s solution (Sigma) at 37°C.

### Immunofluorescence and staining

Oocytes were fixed for 30 min at room temperature with paraformaldehyde 3% in PBS, freshly prepared from a 16% methanol-free paraformaldehyde solution (Electron Microscopy Sciences). Fixed oocytes were permeabilized with 0.1% Triton X-100 (Sigma) in PBS for 15 minutes at room temperature. After blocking with 3% BSA (Sigma) in PBS for 2 h, oocytes were incubated overnight at 4°C with primary antibodies in PBS-BSA 3%. On the next day, oocytes were washed in PBS-BSA 3% and incubated with secondary antibodies diluted 1:1000 in PBS-BSA 1%, for 45 min at 37°C. The following primary antibodies were used : Phospho-myosin light chain 2 (Ser19) (1:200; Cell Signaling Technology #3671), Non-muscle myosin IIA (1:100; Abcam ab24762), Myosin IIB (D8H8) (1:100; Cell Signaling Technology #8824), MRCKα (B-3) (1:100; Santa Cruz Biotechnology sc-374568), MRCKβ (A-2) (1:100; Santa Cruz Biotechnology sc-390127), Tubulin (1:200; Abcam ab6161), p34-ARC (1:100; Santa Cruz Biotechnology sc-515754), Rock-1 (K-18) (1:100; Santa Cruz Biotechnology sc-6056), Trimethyl-histone H3 (Lys4) (1:100; Merck Millipore 04-745). Secondary antibodies were Alexa Fluor 488-conjugated donkey anti-goat, donkey anti-rabbit and goat anti-rat, and Alexa Fluor 555-conjugated goat anti-mouse and goat anti-rabbit (all 1:1000; Invitrogen). Actin filaments were stained with Alexa Fluor 568-phalloidin (Life Technologies). Chromatin was stained with TO-PRO-3 (Invitrogen) or Sir-DNA (Spirochrome).

### Plasmids, cRNA preparation and microinjection

The following plasmids were used: H2B-mCherry in pcDNA3 (Robert Benezra; Addgene plasmid #20972), pGEMHE-eGFP-MAP4 (Jan Ellenberg; Euroscarf plasmid #P30518). MRCKβ-EGFP and MRCKβ (H1593/1596A)-EGFP in pEGFP-N1 were kindly donated by Prof. Shigetomo Fukuhara (Institute of advanced medical science, Tokyo, Japan) and were subcloned into pcDNA3.1. The pEGFP-RhoA Biosensor was from Michael Glotzer (Addgene plasmid #68026) and was subcloned into pcDNA3.1 while replacing EGFP by mCherry. Polyadenylated cRNAs were synthesized in vitro from linearized plasmids, using the mMessage mMachine T7 kit and Poly(A) Tailing kit (Ambion), then purified with RNeasy purification kit (Qiagen), and stored at −80°C. Metaphase II oocytes were injected with ~5pl cRNA solution and cultured for at least 2 h to allow for protein expression.

### Confocal imaging and image processing

For immunofluorescence experiments, oocytes were placed on glass-bottom dishes (MatTek) in a small drop of PBS-BSA 1% supplemented with TO-PRO-3, and covered with mineral oil. Fixed oocytes were imaged with a Leica SP5 or SP8 confocal microscope, using a 63x oil-immersion objective. When appropriate, displayed confocal images were cropped so as to hide excessive non-specific signal arising from the zona pellucida. For live imaging of meiosis resumption and PB2 formation, oocytes were deposited at the center of a glass-bottom dish (MatTek) filled with 2 ml of Ca^2+^-free and BSA-free M16 medium containing 10 mM SrCl2. Time-lapse recordings were aquired using 20x or 40x oil-immersion objectives. The temperature was maintained at 37°C using a stage top incubator (INUBG2E-GSI, Tokai Hit, Shizuoka-ken, Japan) fitted on the microscope stage. Confocal image thickness was set to 1 μm. Confocal images and time-lapse movies were processed with FIJI. Fluorescence intensity profiles were obtained using the segmented line tool and the Plot Profile function in FIJI. Measurements of the chromosome distance to the cortex, and male pronucleus distance to the cortex, were realised using the Leica LAS AF Lite 2.6.0 software.

### Statistical analysis

Statistical analyses were performed using Fisher’s exact test or Student’s t-test, in Origin (OriginLab) or GraphPad QuickCalcs (graphpad.com). Statistical significance was considered when *p*<0.05. Graphics (fluorescence intensity profiles, box plots, scatter plots) were produced in Origin. Box plots show the median (line), 25^th^, 75^th^ percentile (box) and 5^th^, 95^th^ percentile (whiskers).

## Acknowledgements

We thank Prof. Shigetomo Fukuhara (Institute of advanced medical science, Tokyo, Japan) for providing the plasmids encoding MRCKβ-EGFP and MRCKβ (H1593/1596A)-EGFP. We are grateful to the staff of the ARCHE-Biosit animal facility and MRIC-Biosit microscopy facility for technical assistance and expert advice.

## Author contributions

GH conceived and supervised the study. AB, BD and GH designed and performed experiments, and analysed data. GH prepared the figures and wrote the manuscript with input from AB and BD.

## Supplementary Figure legends

**Figure S1.**
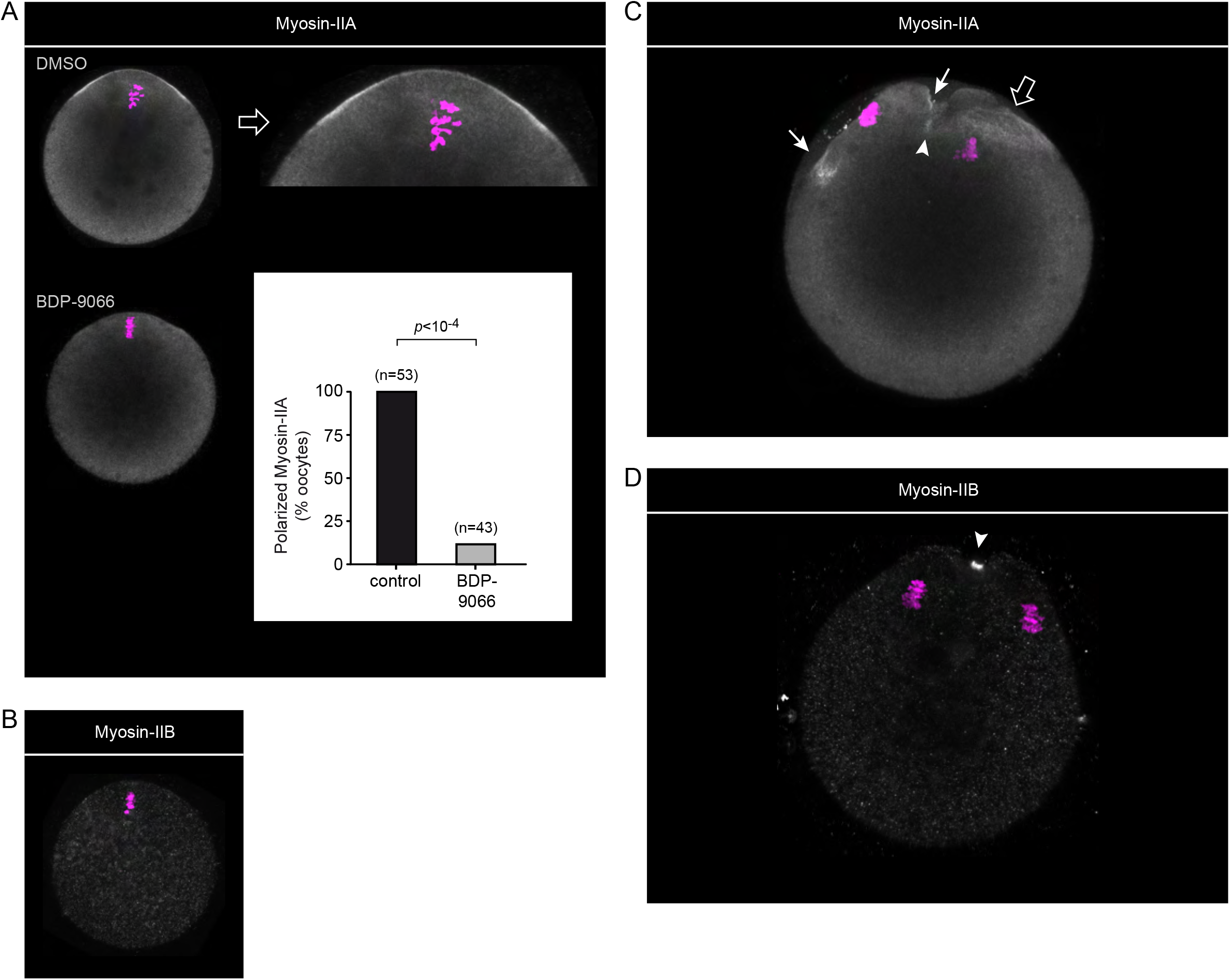
Myosin-IIA and -IIB heavy chain localization in MII and AII oocytes. (A) Immunofluorescence detection of myosin-IIA heavy chain in fixed MII oocytes treated with DMSO (top) or BDP9066 (bottom). An enlarged view of the polarized cortex is shown for the control (DMSO) oocyte. The bar graph displays the fraction of oocytes showing a polarized accumulation of myosin-IIA under each experimental condition. *P* value was calculated using Fisher’s exact test. The number of oocytes scored is indicated above each bar. (B) Immunofluorescence detection of myosin-IIB heavy chain in a fixed MII oocyte. Note the absence of cortical localization. The image is representative of 44 similar observations. (C) Immunofluorescence detection of myosin-IIA heavy chain in a fixed AII oocyte that was activated with SrCl2. The arrows point to the myosin-IIA ring in the nascent polar body. The heavy arrow point to myosin-IIA reorganized into a cap above the internalized chromatid cluster. The white arrowhead point to myosin-IIA in the cleavage furrow. (D) Immunofluorescence detection of myosin-IIB heavy chain in a fixed AII oocyte that was activated with SrCl2. The white arrowhead point to myosin-IIB concentrated in the cleavage furrow. DNA was stained with TO-PRO-3.

**Figure S2.**
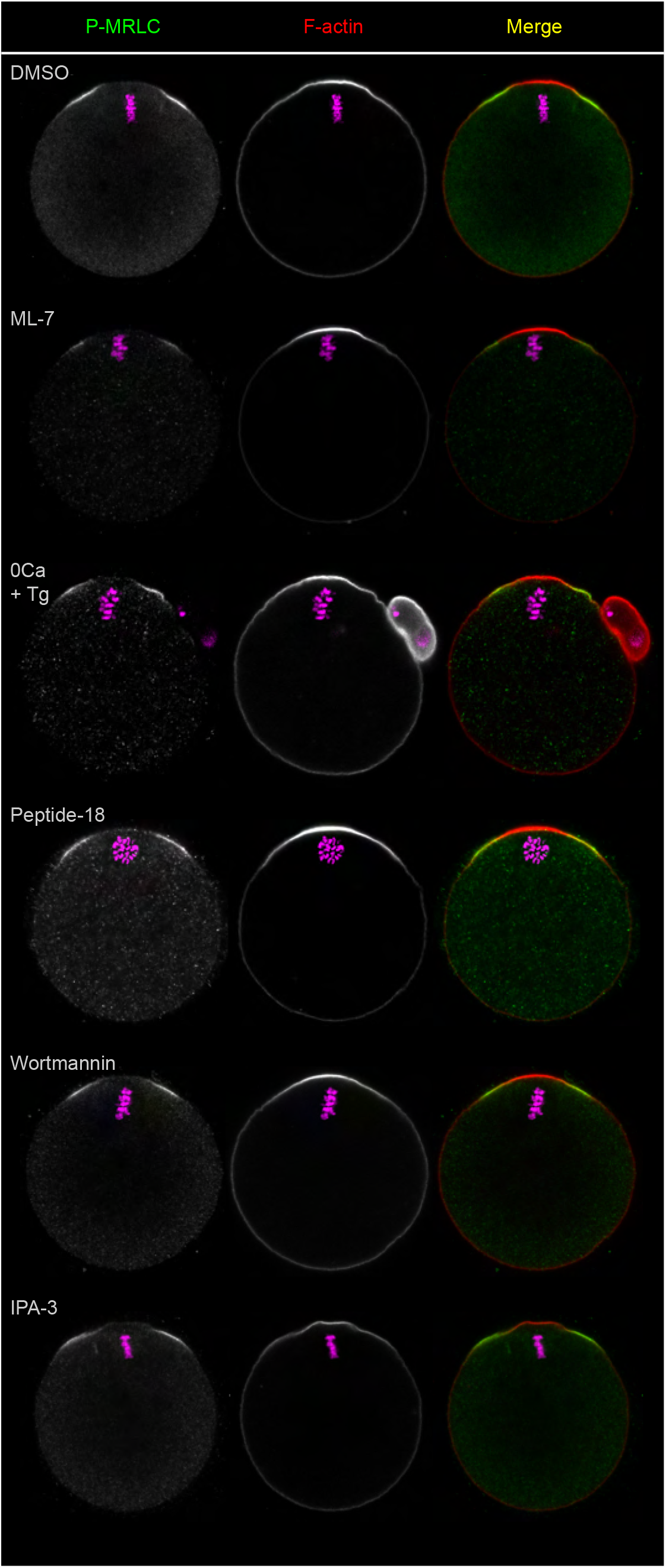
Ring myosin II activation is insensitive to MLCK inhibitors. Ring myosin II activation (P-MRLC) was monitored in MII oocytes after the following treatments (top to bottom): DMSO, ML-7 (15 μM, 1h), Ca^2+^-free medium containing 1 mM EGTA and 10 μM thapsigargin (1h), peptide-18 microinjection followed by a 3-h culture, wortmannin (1 μM, 1h) and IPA-3 (1 μM, 1h). F-actin was labeled with Alexa Fluor 568-phalloidin. DNA was stained with TO-PRO-3.

**Figure S3.**
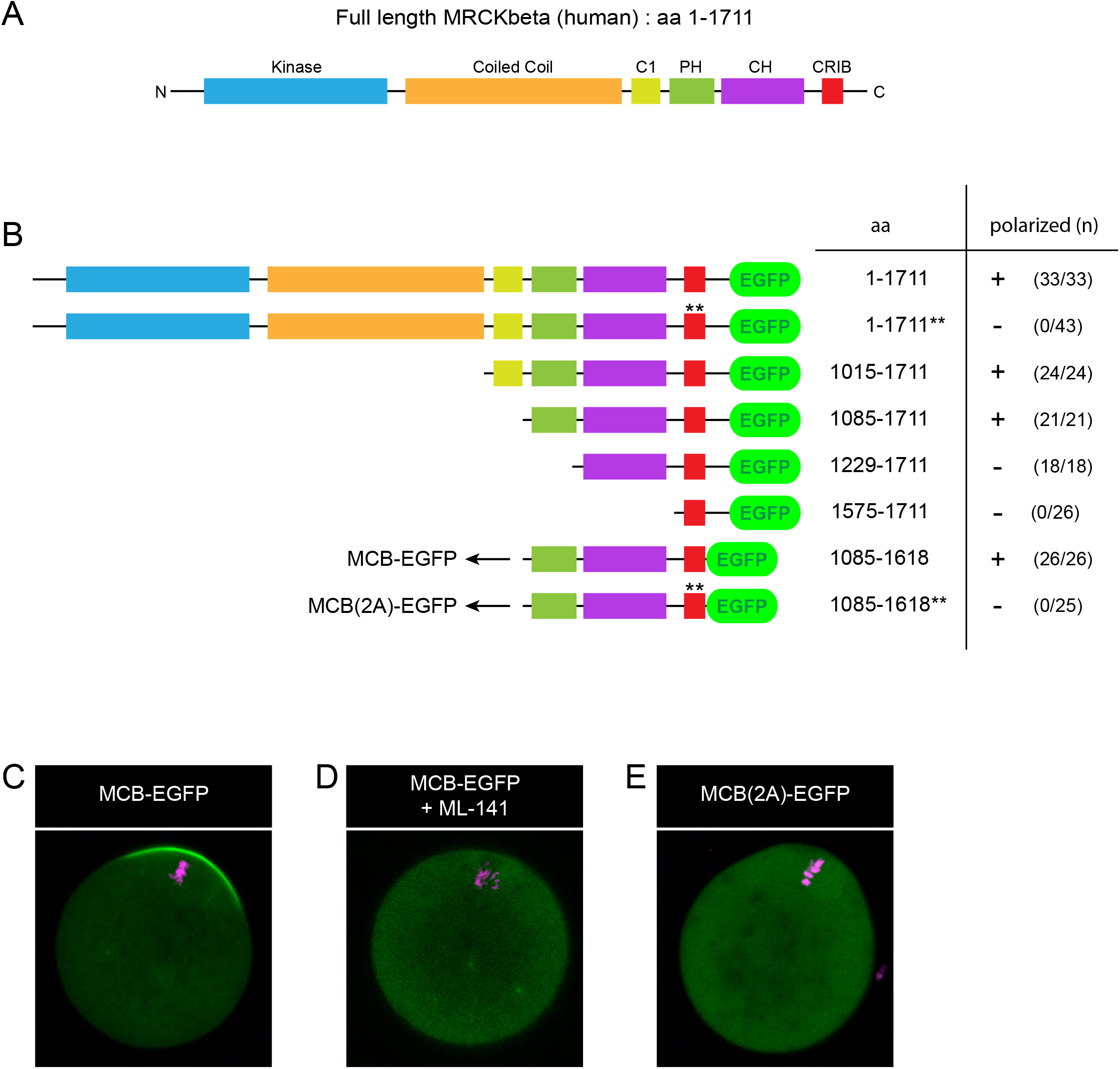
Design of the Cdc42·GTP biosensor MCB-EGFP. **(A)** Schematic depicting the primary structure of full-length human MRCKβ and its functional domains. C1: protein kinase C conserved region 1. PH: pleckstrin homology. CH: citron homology. CRIB: Cdc42/Rac interactive binding. **(B)** Schematic outlining the different constructs generated through stepwise truncation to obtain a Cdc42·GTP biosensor, using full-length MRCKβ-GFP as a template. All constructs were expressed in MII oocytes via cRNA injection, and examined for polarized localization in the cortex overlying maternal chromosomes. The total number of oocytes scored is indicated in parentheses for each construct, with an indication of cortical polarization (+) or absence of (-). While the CRIB domain alone was not sufficient for cortical localization in vivo, stepwise extensions toward the N-terminus allowed us to define a minimal Cdc42·GTP-binding fragment encompassing the PH, CH and CRIB domains. In this study, this minimal fragment was used as a biosensor for Cdc42 activation, designated as MRCK-derived Cdc42·GTP Biosensor (MCB). Mutation of the two key histidine residues in the CRIB domain to alanine, corresponding to H1593/1596A in the full-length sequence, and indicated by a double asterisks (**), abolished polarized localization. aa: amino acids. **(C)** Confocal image of a live MII oocyte expressing the MCB-EGFP biosensor. **(D)** Confocal image of a live MII oocyte expressing the MCB-EGFP biosensor, and treated with ML-141 (5 μM, 1h). **(E)** Confocal image of a live MII oocyte expressing the MCB(2A)-EGFP biosensor, bearing the H1593/1596A substitution. Chromosomes were stained with SiR-DNA.

**Figure S4.**
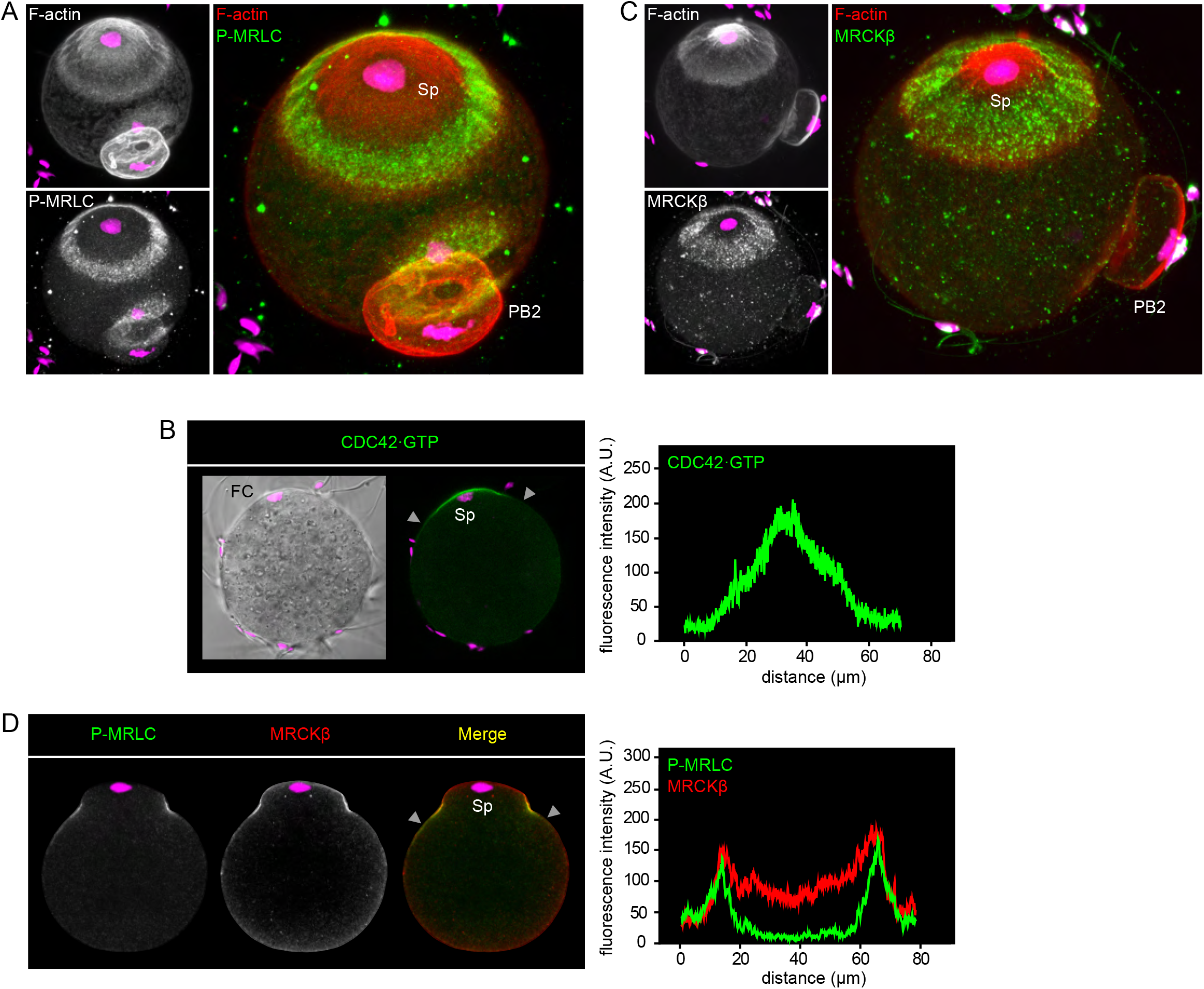
The Cdc42/MRCK/myosin II pathway is activated in the fertilization cone. **(A)** Immunofluorescence detection of activated myosin II (P-MRLC) in a fertilized oocyte. Images are z-compressions across the entire zygote volume, for 3d rendition, highlighting the FC area. **(B)** Detection of Cdc42·GTP in the FC. The image shows a fertilized oocyte expressing the Cdc42·GTP biosensor MCB-EGFP. The oocyte was fertilized in vitro after zona pellucida removal. **(C)** Immunofluorescence detection of MRCKβ in a fertilized oocyte. Images are z-compressions across the entire zygote volume, for 3d rendition, highlighting the FC area. **(D)** Immunofluorescence detection of activated myosin II (P-MRLC) and MRCKβ in a fertilized oocyte. Fluorescence intensity profiles correspond to the cortical regions delineated by the grey arrowheads in the merge images. FC: fertilization cone. Sp: sperm chromatin. PB2: second polar body. F-actin was labeled with Alexa Fluor 568-phalloidin. DNA was stained with TO-PRO-3.

## Supplementary video legends

**Video 1.** Live imaging of anaphase-II in a control oocyte treated with DMSO. The oocyte was injected with cRNAs encoding EGFP-MAP4 to label microtubules and H2B-mcherry to label chromosomes.

**Video 2.** Live imaging of anaphase-II in an oocyte treated with BDP9066, showing the spindle distorsion phenotype. The oocyte was injected with cRNAs encoding EGFP-MAP4 to label microtubules and H2B-mcherry to label chromosomes.

**Video 3.** Live imaging of anaphase-II in an oocyte treated with BDP9066, showing the spindle break phenotype. The oocyte was injected with cRNAs encoding EGFP-MAP4 to label microtubules and H2B-mcherry to label chromosomes.

